# Adgrd1 deficiency reveals increased hippocampal vulnerability and selective behavioral alterations in mice

**DOI:** 10.64898/2026.07.28.741204

**Authors:** Inés Martínez-Soria, Pol Picón-Pagès, Anna P. Pérez- González, Núria Moral, Karolina Zimkowska, Elena de Cecco, Jordi Duran, Xavier Gasull, Adriano Aguzzi, Rosalina Gavín, José Antonio del Río

## Abstract

ADGRD1 (GPR133) is an orphan adhesion G protein-coupled receptor (aGPCR) that primarily signals through G_s_ to regulate intracellular cAMP levels and is increasingly recognized for its roles across multiple tissues, including the central nervous system. Its conserved expression in neural tissues, the presence of splice variants in the fetal brain, and its structural similarity to other aGPCRs all suggest that it might play important roles in the organization of neural circuits. Recent studies have identified a wide range of extracellular, membrane-associated, and intracellular interacting partners, highlighting the receptor’s ability to integrate diverse signals. In this study, we explored the consequences of Adgrd1 deficiency in mice using behavioral, electrophysiological, and transcriptomic approaches. Adgrd1-null mice showed reduced nest-building behavior and decreased exploratory drive, while motor coordination and recognition memory remained largely intact. Electrophysiological recordings indicated a trend toward impaired long-term potentiation. These mice also exhibit increased susceptibility to kainate-induced excitotoxicity. RNA-seq analysis revealed coordinated changes in genes associated with inhibitory signaling, extracellular matrix organization, and cytoskeletal regulation, pointing to a shift toward reduced synaptic stabilization and increased hippocampal vulnerability. Altogether, these results suggest that Adgrd1 plays a key role in maintaining hippocampal resilience and regulating motivational behaviors through integrated molecular and circuit-level mechanisms.

## Introduction

The adhesion G protein-coupled receptor ADGRD1 (also known as GPR133) is an orphan adhesion G protein–coupled receptor (aGPCR) that couples predominantly to G_s_ proteins, thereby modulating intracellular cAMP production in several cell types ^1–4^. ADGRD1 has gained attention as a multifunctional aGPCR with emerging relevance in several human tissues (please see https://gtexportal.org/home/gene/ADGRD1), including the central nervous system in healthy and disease conditions (e.g. ^5^). Its expression in both human and mouse brain tissue was first documented through Expressed Sequence Tag (EST) analyses, indicating a conserved neural presence ^6^. Subsequent work identified multiple splice variants in fetal brain, a finding that appears to indicate that ADGRD1 could participate in neurodevelopmental programs ^7^. Although the precise mechanisms remain fully determined, structural and functional studies of adhesion GPCRs suggest that receptors in this family, including ADGRD1, might contribute to processes such as neural progenitor proliferation, cortical organization, and developmental signaling^8,9^. Additional evidence from glioblastoma research shows that ADGRD1 is active in human brain-derived cells and responsive to hypoxic cues, further supporting its physiological relevance within CNS environments; indeed, its expression is required for tumor growth under hypoxic conditions, where it functions as a marker of malignancy and is transcriptionally upregulated by Hypoxia-Inducible Factor 1 alpha (HIF1α) _10,11_.

In this regard several extracellular ligands, membrane-associated co-receptors, and intracellular binding partners have been identified for ADGRD1. Recent studies have reported multiple molecules capable of interacting with the receptor across distinct cellular contexts, including the extracellular androgen hormone 5α-dihydrotestosterone (5α-DHT) ^12^, the transmembrane coreceptor PTK7 ^13^, the extracellular matrix-associated Plexin Domain-Containing Protein 2 (PLXDC2) ^13^, the intracellular lipid-transfer protein Extended Synaptotagmin-1 (ESYT1) ^14^ or the small molecule GL64 ^15^. Collectively, these findings underscore the molecular versatility of ADGRD1 and suggest that its activation integrates signals from soluble ligands, membrane-embedded partners, and intracellular effectors.

Somewhat independently of those functional studies, A. Aguzzi’s laboratory carried out, ten years ago, a pioneering systematic screen to identify GPCRs that can interact with the flexible N-terminal domain (FT) of the non-pathogenic cellular prion protein (PrP^C^) ^16^. As a result of this study, the laboratory described in detail the functions of GPR126 (Adgrg6), during peripheral myelinization concluding that PrP^C^ promotes myelin homeostasis through FT-mediated GPR126 agonism ^16^. In the same study, two members of the superfamily were described as having the opposite effect. When HEK293 cells expressing ADGRG1 (GPR56), but especially ADGRD1, were treated with the N-terminal fragment (23-50) of PrP^C^, intracellular cAMP levels decreased in the treated cells _16_, reviewed in ^17^. In the absence of direct molecular confirmation, the functional data allow us to propose that ADGRD1 and ADGRG1 could share some parallel signaling roles with PrP^C^.

Numerous functions and processes have been ascribed to PrP^C^, implicating it in roles that span neural development to neurodegeneration, including effects on behavior ^18–27^. Deficiency or dysfunction of PrP^C^ has been linked to mood disorders, including depression and bipolar disorder, in both animal models and human studies (e.g., ^28–30^). On the other hand, we and others demonstrated both *in vitro* and *in vivo* that the absence of PrP^C^ leads to increased susceptibility to kainate ^26,31–38^. Indeed, different PrP^C^-deficient animal models with different genetic background consistently exhibit this increased susceptibility and display impairments in memory and behavior associated with the absence of this protein _26,31_. With these considerations in mind, the present study aims to initiate the characterization of the behavioral and physiological phenotype associated with Adgrd1 deficiency in a mouse model, with a particular focus on determining whether the absence of this receptor influences both seizure susceptibility responses to kainate and behavior in young-adult mice.

Results indicate that Adgrd1 mutant mice show increased susceptibility to kainate and reduced naturalistic behavior, evidenced by impaired nest building indicating preserved species-typical repetitive behaviors. In the open field, mutants spend more time in the center but exhibit markedly reduced locomotion, limiting standard anxiety-related interpretations and instead suggesting diminished exploratory drive or general activity. Electrophysiological analyses reveal a trend toward reduced hippocampal LTP and alterations in the GABA_A_ receptor α2 subunit, consistent with heightened hippocampal vulnerability. Motor coordination is preserved, and although novel object recognition performance remains largely intact, reduced object interaction suggests deficits in motivation or exploratory drive rather than impaired recognition memory. Overall, the phenotype is consistent with hippocampal vulnerability, accompanied by selective alterations in motivational and affective domains, while core motor and basic cognitive functions remain largely preserved. Notably, this profile differs from that described in PrP^C^-deficient mice, suggesting that Adgrd1 may exert pleiotropic roles in the brain, potentially involving multiple yet unidentified ligands. As Adgrd1 is also expressed in peripheral tissues, we cannot exclude the contribution of peripheral alterations to the observed behavioral and electrophysiological phenotypes. In particular, systemic changes in metabolic, endocrine, or immune signaling may influence neuronal excitability and behavioral output.

## Materials and Methods

### Animals

Adult (2-7 months) C57BL/6J mice (*Adgrd1^+/+^*) were purchased from Charles River Laboratories (Paris, France). *Adgrd1^0/0^* mouse strain was provided by A. Aguzzi. A total of 89 adult mice were included in this study, with 44 *Adgrd1^0/0^* (13 females and 31 males) and 45 *Adgrd1^+/+^*(14 females and 31 males) mice. All procedures were conducted in accordance with the protocols and guidelines set by the Ethical Committee for Animal Experimentation (CEEA) of the University of Barcelona. CEEA approved the protocols for working with animals in this study (CEEA approval 214/24, OB 235/24, OB 41/21 and C/EB517.22). Behavioral and electrophysiological experiments were performed in compliance with the European Union Council guidelines (2010/63/EU) and current Spanish legislation (BOE 34:11370-421, 2013) for the use of laboratory animals in chronic experiments.

### *Adgrd1^0/0^* mice validation by PCR

Primers sequences for *Adgrd1^0/0^* genotyping were provided by A. Aguzzi. The sequences were: Forward: 5′-CTGGGGTGAGAGATGTGCTG-3′ and Reverse: 5′-CTACCCACTCCTCTACCAGCC-3′. The PCR protocol was: 95 °C (5 min), 35 cycles of: 95°C (45 s), 60°C (45 s), 72°C (45 s), and 72°C (2 min). Samples were kept at 4°C until removed from the thermocycler. For *Adgrd1^0/0^* animals, the expected size of the PCR product on the electrophoresis gel was 170 bp (Supplementary Figure 1A).

### Behavioral studies

A total of 62 mice were included in these experiments, comprising of 16 male mice *Adgrd1^+/+^*, 19 male mice *Adgrd1^0/0^*, 14 female mice *Adgrd1^+/+^* and 13 female mice *Adgrd1^0/0^*. Mice were kept in cages with a 12-hour light/dark cycle, maintained at a consistent room temperature of 21 ± 1°C. Both food and water were provided *ad libitum*.

### Nest building

For this experiment, 7 *Adgrd1^+/+^* and 9 *Adgrd1^0/0^* male mice and 4 *Adgrd1^+/+^* and 3 *Adgrd1^0/0^* female mice were used. On the first day of the experiment, two square cotton pieces (5 × 5 cm) were placed in each cage to encourage nest building. Nest presence and quality were documented by photographs and evaluated the following day using a modified 5-point scale (0-4), as described by ^39^. Two researchers, blinded to group allocation, independently assessed the nests.

### Limb Clasping Test

For this experiment, 9 *Adgrd1^+/+^* male mice and 10 *Adgrd1^0/0^*male mice, as well as 10 female mice of each genotype, were used. The procedure followed Miedel *et al*., ^40^, involving tail suspension for 5-10 seconds while recording video, ensuring both the fore paws and hind paws were visible during suspension to observe limb clasping behavior. Videos were reviewed, and limb clasping was scored on a 0-4 scale. Two blind investigators scored the mice, and if their scores differed by more than one point, the mouse was rescored, with the final score being the average of both investigators.

### Open-field test

In this study, the locomotor activity of mice was tested in a white square open-field arena (50 × 50 × 40 cm). 7 animals of each sex and genotype, a total of 28 mice, were used. Mice were placed in the center of the arena for 20 minutes, and their locomotor activity was tracked and analyzed using MouBeAT software ^41^. Mice behavior was evaluated by designating the central square of the arena (30 × 30 cm) as the center zone. Time spent in the peripheral zone was used to assess thigmotaxis, an indicator of anxiety-related behavior ^42^. Additionally, fecal pellets were counted, as increased defecation often reflects heightened emotional responses ^42^. The arena and walls were disinfected with 70% ethanol between each trial.

### Novel object recognition test

The object recognition test was conducted using the same arena as the open field test. 7 animals of each sex and genotype, a total of 28 mice, were used. The experiment consisted of three phases, each lasting 20 minutes. In the first phase, mice explored the empty arena. On the second phase, two identical objects were placed diagonally in the box. On the final phase, one object was replaced with a different one in shape and color. The arena and objects were disinfected with 70% ethanol between trials. Mice behavior was recorded and analyzed using MouBeAT software ^41^, with exploration defined as head approaches or touches to the objects.

### Kainate-induced epilepsy and seizure analysis

A total of 16 *Adgrd1^0/0^* and 17 *Adgrd1^+/+^*male mice were used in these experiments. This study was carried out using as a reference the procedure previously published by our group ^31^. A kainate (KA) solution (Sigma-Aldrich, Darmstadt, Germany) dissolved in 0.1M phosphate buffered saline (PBS, pH 7.35) was administered in three injections (8 mg/kg body weight (b.w.) at 0, 30, and 60 minutes). After the first injection, animals were housed in clean boxes with two mice per box. The behavior of the treated mice and the occurrence of seizures were monitored and video-recorded for 4 hours. Seizure severity was classified into grades: grades I-II: reduced activity and immobility; grades III-IV: increased activity and scratching behavior; grade V: impaired balance and sporadic convulsions; grade VI: continuous seizures characterized by bouncing or “popcorn-like” behavior, which may include blinking episodes ^31^.

### Fluoro-Jade B staining

The following day or five days after KA administration, mice were perfused with 4% phosphate buffered paraformaldehyde (PFA, pH 7.35). Brains were dissected and postfixed overnight (O/N), then cryoprotected in PBS containing 30% sucrose (w/v). After freezing in dry ice, 30 μm coronal sections were cut using a freezing microtome (Leica Microsystems, Wetzlar, Germany). Coronal sections containing the dorsal hippocampus were selected, rinsed in 0.1M Tris-HCl pH 7.4 buffer and mounted on gelatin-coated slides. These slides were then dried at 37°C O/N. Sections were pretreated with ethanol, oxidized with 0.06% KMnO_4_ (Sigma-Aldrich) and incubated with 0.001% Fluoro-Jade B (Biosensis Pty Limited, Davis, CA) and 0.01% DAPI (Thermo Fisher Scientific, Waltham, MA) in 0.1% acetic acid. After rinsing, clearing and drying; sections were coverslipped with Eukitt^TM^ (ITW Reagents Panreac, Barcelona, Spain). Fluoro-Jade B fluorescence in the hippocampus (4 sections of each mouse, n = 3 mice per genotype and treatment) was documented using an Olympus BX61 epifluorescence microscope with a cooled DP72L camera (Olympus, Tokyo, Japan). Photomicrographs were taken with identical time exposure between preparations from each wild-type and respective *Adgrd1^0/0^* mouse.

### Immunohistochemical procedures

Mice were perfused with 4% PFA (pH 7.35), and their brains were dissected and postfixed O/N before being cryoprotected in PBS containing 30% sucrose (w/v). After freezing, 50 μm coronal sections were cut using a freezing microtome. For immunohistochemistry, free floating brain sections were treated to block endogenous peroxidase activity by incubating in 3% hydrogen peroxide (H_2_O_2_) and 10% methanol in PBS, followed by incubation in a blocking solution for 2-4 hours at room temperature (RT). Sections were then incubated with primary antibodies O/N at 4°C (see Table 1). For bright-field microscopy, species-specific biotinylated secondary antibodies were applied for 2 hours at RT (see Table 1). An avidin-biotin-peroxidase complex (ABC) was applied following the manufacturer’s protocol (Vector Laboratories, Newark, CA), and peroxidase activity was visualized using 0.03% 3,3’-diaminobenzidine (DAB) and 0.01% H_2_O_2_, in some cases together with ammonium nickel sulfate or cobalt chloride to enhance signal contrast ^43,44^. For immunofluorescence, secondary antibodies (Alexa Fluor 488 or 568) were applied for 2 hours at RT, followed by incubation with 1 μg/mL DAPI in PBS for 10 minutes at RT, and samples were then mounted in Mowiol™ (EMD Millipore, Burlington, MA). Labeled sections (n = 3-4 mice, 3- 15 slices per mouse) were imaged using an Olympus BX61 microscope with a cooled digital DP72L camera (Olympus, Tokyo, Japan). Image processing was done using ImageJ™ software.

**Table 1.** Antibodies used in the present study.

| **Antibody** | **Host** | **Catalog number** | **Supplier** | **Dilution** | **Application** |
| --- | --- | --- | --- | --- | --- |
| **Phospho-p44/42 MAPK (Erk1/2) (Thr202/Tyr204)** | Rabbit | 9101S | Cell Signaling Technology (CST) | 1:100 | ICC |
| **Anti-Glial Fibrillary Acidic Protein (GFAP)** | Mouse | MAB360 | Chemicon | 1:100 | ICC |
| **Adgrd1** | Rabbit | DF4947 | Affinity Biosciences | 1:100 | ICC |
| **Iba1** | Guinea Pig | 234004<br>(discontinued) | Synaptic Systems | 1:125 | ICC |
| **Goat anti-Guinea Pig IgG (H+L) Secondary Antibody, Alexa Fluor 568** | Goat | A-11075 | Invitrogen (Thermo Fisher Scientific) | 1:1000 | ICC |
| **Biotinylated anti-Rabbit IgG (H+L)** | Goat | BA-1000 | Vector Laboratories | 1:150 | ICC |
| **Goat anti-Rabbit IgG (H+L) Secondary Antibody, Alexa Fluor 568** | Goat | A-11011 | Invitrogen (Thermo Fisher Scientific) | 1:1000 | ICC |
| **Goat anti-Mouse IgG (H+L) Secondary Antibody, Alexa Fluor 488 conjugate** | Goat | A-11029 | Invitrogen (Thermo Fisher Scientific) | 1:1000 | ICC |

### Electrophysiological recordings in hippocampal slices

A total of 14 mice (7 *Adgrd1^0/0^* and 7 *Adgrd1^+/+^*) were used in these experiments. Animals were anesthetized using isoflurane and decapitated. To obtain brain slices, this organ was rapidly removed and placed in ice-cold artificial CSF1 (aCSF1, in mM; 206 sucrose, 1.25 NaH_2_PO_4_, 26 NaHCO_3_, 1.3 KCl, 1 CaCl_2_, 10 MgSO_4_, 11 glucose, purged with 95% O_2_ / 5% CO_2_, pH: 7.35). After removing olfactory bulbs and cerebellum, coronal slices (380-400 µm) were obtained with a vibratome (VT1200S; Leica Microsystems) in the same cold solution. Slices were placed in an incubator beaker containing artificial CSF2 (in mM, 119 NaCl, 1.25 NaH_2_PO_4_, 25 NaHCO_3_, 2.5 KCl, 2.5 CaCl_2_, 1.5 MgSO_4_, 11 glucose, purged with 95% O_2_ / 5%CO_2_, pH: 7.35). Slices were kept at 34°C for 1h and subsequently at RT for at least one additional hour before starting the recordings. Slices were transferred into a measurement chamber continuously perfused with carbogenated aCSF2 at 27°C for field excitatory post-synaptic potential (fEPSP) recordings. A bipolar stimulation electrode was placed in the Schaffer’s collateral pathway of the hippocampus. The recording electrode filled with aCSF2 was placed in the dendritic branching of the CA1 region. A stimulus isolation unit A385 (WPI, Hertfordshire, UK) was used to elicit stimulation currents between 25-180 µA. Before baseline recordings, input-output (IO) curves were recorded for each slice at 0.03 Hz. The stimulation current was then adjusted in each recording to evoke fEPSPs at which the slope was at 50-60 % of maximally evoked fEPSP slope value. After baseline recording for 15-20 minutes at 0.03 Hz; long-term potentiation (LTP) was induced by theta-burst stimulation (TBS; 7 trains with 30 s intervals, with each train containing 10 bursts at 5 Hz and each burst containing 4 pulses at 100 Hz). After LTP induction, fEPSPs were recorded for one additional hour at 0.03 Hz. Short-term plasticity was evaluated as paired-pulse facilitation. Two consecutive fEPSPs were measured before LTP induction at 0.03 Hz with an interstimulus interval of 50 or 100 ms. The facilitation was assessed as the ratio between the first and second stimuli. All recordings were amplified and stored using amplifier AxoClamp 2B (Molecular Devices, San Jose, CA). Traces were analyzed using Axon pClamp software (Molecular Devices, version 10.6).

### Bulk RNAseq analysis

Total hippocampal RNA was extracted and processed by Haplox (Shenzhen, China). RNA purity was assessed using NanoDrop™ One/OneC spectrophotometry (Thermo Fisher Scientific), and concentration was quantified with a Qubit™ RNA HS Assay (Thermo Fisher Scientific). RNA integrity was evaluated on an Agilent 4200 TapeStation (4200 TapeStation, Agilent Technologies, Santa Clara, CA) and only samples with RIN ≥ 7.0 were accepted for library preparation. Poly(A)+ mRNA was isolated using oligo-dT magnetic beads, fragmented, and reverse-transcribed to synthesize first-strand cDNA using M-MuLV reverse transcriptase, followed by second-strand synthesis with DNA polymerase I. After end repair, A-tailing, adaptor ligation, and size selection (∼200 bp) with AMPure XP beads, libraries were amplified by PCR and quality-checked by qPCR (KAPA) and TapeStation. Sequencing was performed on an Illumina platform (PE150, Illumina Inc., San Diego, CA). Raw reads were processed with *fastp* for adapter trimming, removal of low-quality bases, and sliding-window filtering ^45^. Clean reads were aligned to the mouse reference genome using HISAT2 ^46^, and BAM files were sorted and filtered with SAMtools_47_. Gene-level quantification was performed with HTSeq (union mode) ^48^.

Differential expression analysis was conducted using DESeq2 with Benjamini– Hochberg correction, applying |log2FC| ≥ 1 and padj < 0.05 as significance thresholds ^49^.

### RT-qPCR assays

Total RNA from hippocampal tissues was isolated using the mirVana RNA isolation kit (Thermo Fisher Scientific) according to the manufacturer’s protocol. RNA quantity was determined using a Qubit fluorometer, and RIN values were assessed by the Genomics Service of the Scientific and Technological Services at the University of Barcelona (CCiTUB). cDNA for was obtained using the SuperScript™ II Reverse Transcriptase kit (Thermo Fisher Scientific) following the supplier’s guidelines. Each RT-qPCR reaction was done using LightCycler® 480 SYBR Green I Master kit and in triplicate using a 96-well optical plate (Roche Diagnostics, Mannheim, Germany). The reaction was done with ≍45 ng of cDNA, and previously used primers (Table 2). PCR amplification and melt curve were conducted using the StepOnePlus Real Time PCR System (Applied Biosystems, Foster City, CA). The relation between expression of housekeeping gene Hypoxanthine Phosphoribosyltransferase 1 (HPRT) was used, outliers were discarded based on both amplification and melt curves. Data were analyzed following the 2^-ΔCt^ method.

**Table 2.**
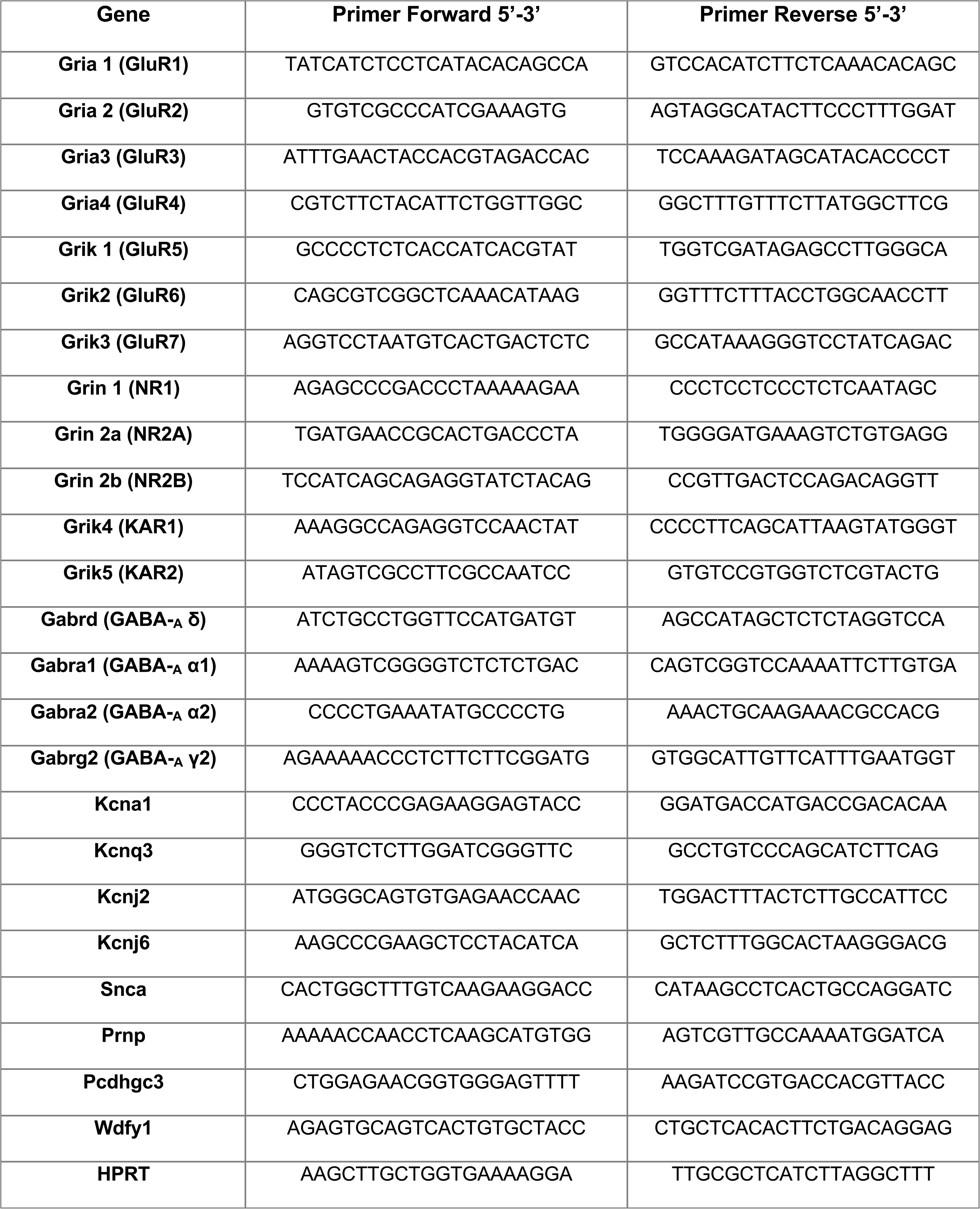
Primers used in RT-qPCR experiments.

### Statistical analysis

Data analyses were carried out using GraphPad Prism versions 10.0.1.218 (ID: 97F7835C03E). Normality of the data distributions was evaluated using the Shapiro-Wilk test. Outliers in behavioral experiments were identified using the ROUT method (robust regression and outlier removal) implemented in GraphPad Prism, with a false discovery rate (Q) of 10%. Statistical analyses were performed using parametric or non-parametric tests, depending on the data distribution and experimental design. Differences between two groups were assessed using an unpaired Student’s t-test or the Mann-Whitney *U* test, while comparisons among multiple groups (≥ 4) were analyzed by one-way ANOVA followed by Holm-Šídák multiple comparisons test and Multiple unpaired Student’s t-test. Statistical significance was defined as \**p* < 0.05, \*\**p* < 0.01 and \*\*\**p* < 0.001. Results are presented as mean ± standard deviation of the mean (SEM).

## Results

### Immunolocalization of Adgrd1 in the adult mouse hippocampus

In the present study, we tested several antibodies against Adgrd1, most of which failed to immunostain the mouse hippocampus in frozen sections (e.g., LS- C747720, LSBio), even when signal amplification systems using nickel or cobalt enhancement were applied. In contrast, the antibody clone DF4947 (Affinity Biosciences) against Adgrd1 specifically immunodetected the protein in paraffin sections of the mouse renal cortex as a control, displaying a focal and heterogeneous distribution predominantly localized to interstitial cells within the peritubular compartment, whereas glomerular and tubular epithelial structures showed minimal or absent staining (Supplementary Figure 1B). This pattern differs from that reported in human kidney, where tubular structures are also immunopositive (www.proteinatlas.org/ENSG00000111452ADGRD1).

Moreover, this distribution is consistent with publicly available single-cell transcriptomic datasets showing enrichment of Adgrd1 in fibroblast-associated and interstitial cell clusters (https://cellxgene.cziscience.com/gene-expression).

In parallel, when this antibody was used in wild-type adult hippocampal sections following antigen retrieval, we observed immunoreactivity in the pyramidal layer of CA3 closest to the dentate gyrus (Supplementary Figure 1C, D). This staining gradually diminished in pyramidal layer approaching the CA3-CA1 transition (Supplementary Figure 1C, D). In CA1, the signal was very weak and nearly undetectable by immunohistochemistry. Some cells present in the plexiform layers were also slightly stained. In sections developed with DAB enhanced with cobalt or nickel, complete neuronal labelling was observed in the aforementioned regions, along with a small number of labelled cells in plexiform layers (Supplementary Figure 1E). Nickel enhancement further increases the visibility of these sparsely distributed neurons in the plexiform strata, although at the cost of elevating background staining (Supplementary Figure 1E), but CA1 pyramidal cells were not strongly detected. This labeling was not observed in sections from the *Adgrd1^0/0^* mice (Supplementary Figure 1F) and wild-type sections without the primary antibody prevented immunostaining (Supplementary Figure 1G).

### *Adgrd1^0/0^* mice show reduced nesting and exploratory behavior with preserved motor function and sex-specific differences in recognition memory

Next, we examined the behavior of *Adgrd1^0/0^* to obtain a broad characterization of the phenotype, providing an initial overview of potential functional alterations. (Figure 1). To begin with, we performed the nest building test (Figure 1A). The scoring distribution differed markedly between groups with no sex-specific differences observed. *Adgrd1^+/+^*mice showed a bimodal profile, with most animals scoring 1 or 4 (33.33%), and fewer individuals at intermediate values. *Adgrd1^0/0^* mice displayed a compressed distribution centered on score 2 (58.33%), with additional representation at scores 1.5 (8.33%), 2.5 (16.67%) and 3.5 (8.33%), and no animals reaching the maximum nest building score. Group comparison therefore indicates a shift from broad, polarized distribution in controls to a predominantly mid-range scoring pattern in *Adgrd1^0/0^* mice (Figure 1A). In conclusion, *Adgrd1^0/0^* mice show a constrained, mid-range nest building performance, lacking the high scores observed in controls, which display a broader and more polarized distribution. These mice fail to achieve high nest building scores, indicating a reduced motivational drive to perform this goal-directed behavior and suggesting that Adgrd1 may play a role in the neural mechanisms underlying motivation.

**Figure 1.**
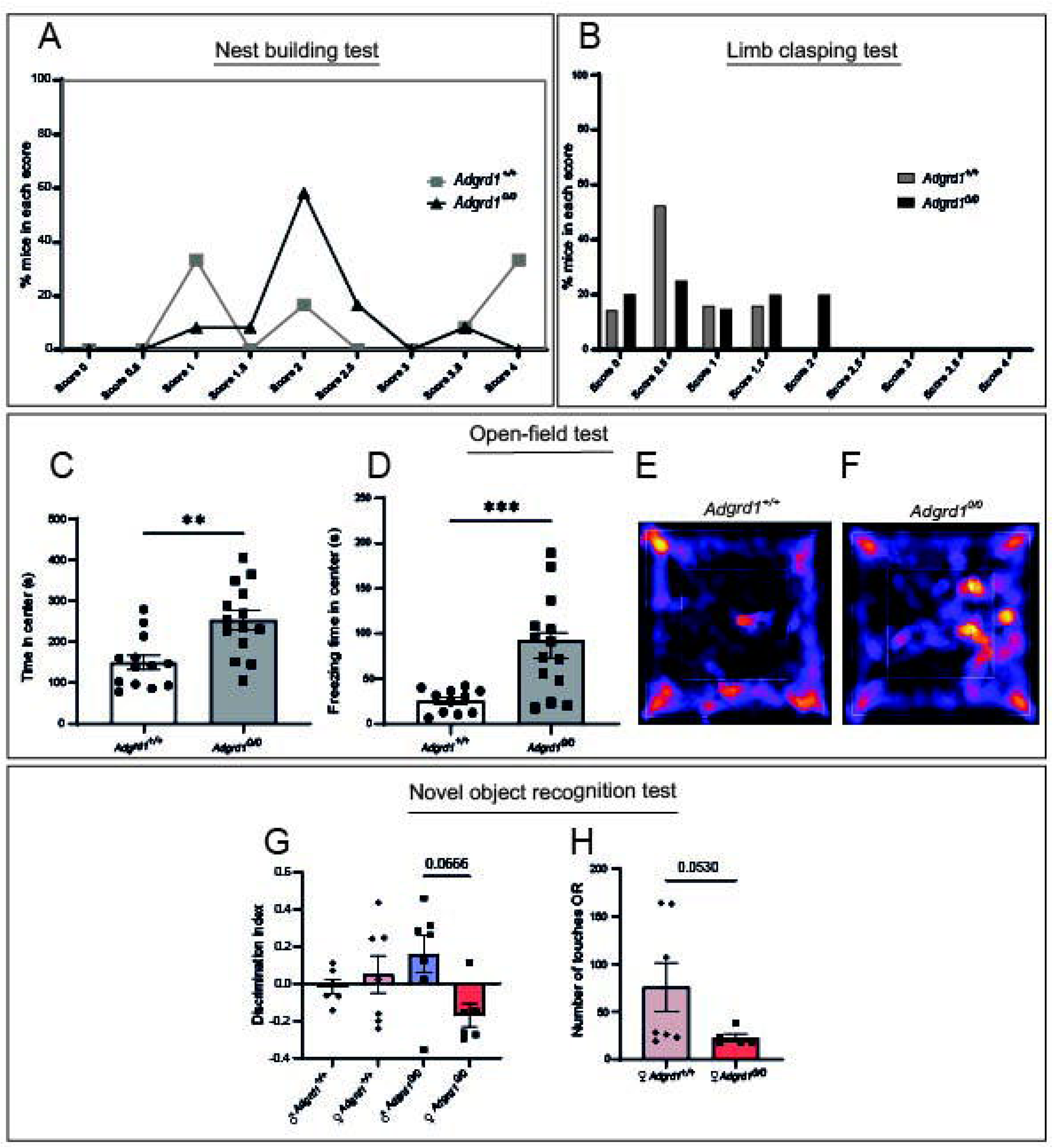
Behavioral characterization of *Adgrd1^0/0^* mice. **A)** Graph illustrating the percentage of *Adrgd1^0/0^* and wild-type mice in the different scores during the nest building test after 20 hours. **B)** Graph showing the percentage of wild-type and mutant mice in the different scores of the limb clasping test. **C)** Time spent in the center of the open-field arena of *Adrgd1^0/0^* and wild-type mice (unpaired t-test: *p* = 0.0018). **D)** Freezing time in the center of open-field arena of *Adrgd1 ^0/0^* and wild-type mice (unpaired t-test: *p* = 0.0005). **E-F)** Heat maps generated by the MouBeAT software from the open-field test illustrating prolonged immobility (red-yellow) in the center of the arena (inner square) in *Adgrd1^0/0^*mice *vs.* wild-type. **G)** Discrimination index in the novel-object recognition test between genotypes and sex (Holm-Šídák’s post-hoc: *p* = 0.0666). **H)** Total number of object touches on day one (OR) in *Adgrd1^0/0^*mice *vs.* wild-type females (Mann-Whitney *U* test: *p* = 0.053).

To assess whether the previously observed mid-range scoring reflected a potential deficit in motor function, we performed the limb clasping test, which showed comparable low-score distributions in both genotypes (Figure 1B). *Adgrd1^+/+^*mice scored within the 0-1.5 range (100 %), with most animals at 0.5 (52.6 %) and no individuals reaching higher values. *Adgrd1^0/0^* mice displayed a similar low-score profile, with most animals also falling between 0 and 1.5 (80 %), and only a 20 % reaching score 2. Group comparison indicated no significant differences between genotypes (Mann-Whitney *U* test: *p* = 0.182; n = 19 *Adgrd1^+/+^* and 20 *Adgrd1^0/0^*), indicating no detectable impairment in limb clasping behavior and suggesting preserved motor function at this level of assessment.

Exploratory behavior was next assessed using the open-field test (Figure 1C-F). *Adgrd1^0/0^* mice spent more time in the center of the arena than *Adgrd1^+/+^* (Figure 1C; unpaired t-test: *p* = 0.0018, n = 13 *Adgrd1^+/+^* and n = 14 *Adgrd1^0/0^*), a pattern that could be interpreted as reduced anxiety. Furthermore, fecal pellets counts did not differ significantly between genotypes (data not shown). However, heat maps and behavioral quantification of freezing time in center revealed that KO animals remained immobile in the center rather than engaging in active exploration (unpaired t-test: *p* = 0.0005, n = 11 *Adgrd1^+/+^* and n = 14 *Adgrd1^0/0^*) (Figure 1D-F). No sex-dependent differences were detected. Although *Adgrd1^0/0^* mice spent more time in the center, this was primarily driven by increased immobility rather than active exploration (Figure 1E, F). Given that animals are initially placed in the center, this behavior is more consistent with reduced exploratory drive or decreased initiation of movement rather than reflecting an anxiolytic-like phenotype.

To further evaluate exploratory motivation, we conducted the novel object recognition test (Figure 1G, H). No significant genotype-dependent differences in discrimination index were observed, with male and female *Adgrd1^+/+^* and *Adgrd1^0/0^* groups showing broadly overlapping distributions. However, a trend toward higher discrimination indices in *Adgrd1^0/0^* males compared to females was detected (Holm-Šídák’s post-hoc: *p* = 0.0666), suggesting a possible sex-dependent effect that did not reach statistical significance. Notably, female *Adgrd1^0/0^* mice exhibited reduced object exploration during the familiarization phase compared to female *Adgrd1^+/+^*, as indicated by fewer object touches when both objects were identical (Mann-Whitney *U* test: *p* = 0.053; n = 7 *Adgrd1^+/+^* and 5 *Adgrd1^0/0^*). This reduced interaction may have contributed to the lower discrimination performance. Overall, these findings are consistent with diminished exploratory engagement rather than a primary impairment in recognition memory.

### The absence of Adgrd1 preserves basal synaptic transmission but impairs LTP maintenance

To further investigate whether the absence of Adgrd1 alters basal synaptic function in the hippocampus, we next analyzed several parameters of basal synaptic activity in acute hippocampal slices obtained from mutant and wild-type animals (Figure 2). Baseline synaptic transmission was indistinguishable between genotypes, as input-output curves from wild-type and *Adgrd1^0/0^* slices showed nearly overlapping fEPSP amplitudes across stimulation intensities, indicating preserved CA3-CA1 synaptic gain. Paired-pulse facilitation (as a form of short-term plasticity) was similar between genotypes, indicating that presynaptic mechanisms were not significantly altered (not shown). Following high-frequency stimulation, both groups displayed clear LTP induction; however, *Adgrd1^+/+^* slices reached higher potentiation levels and maintained a more stable plateau, whereas *Adgrd1^0/0^*slices exhibited weaker and more variable potentiation, particularly during the late phase (Figure 2A). This divergence was evident in the final 10-minute window (50-60 min), where *Adgrd1^+/+^* responses remained consistently elevated (136-180% of baseline), while *Adgrd1^0/0^* values ranged more widely (85-165%), including markedly reduced traces in some recordings. Although the late-phase difference did not reach conventional statistical significance (*p* ≈ 0.057), it showed a clear trend toward impaired LTP maintenance in the knockout group. Quantification at the 10-minute time point further supported this pattern. *Adgrd1^+/+^* slices exhibited exhibited robust potentiation (150.37 ± 6.36%), whereas *Adgrd1^0/0^* slices showed a reduced and more variable response (126.87 ± 9.90%). An unpaired Student’s t-test indicated a trend toward decreased early maintenance in the knockout (*p* = 0.0691), though not statistically significant (Figure 2B). Together, these findings indicate that loss of Adgrd1 does not affect basal synaptic transmission or LTP induction but selectively compromises the stabilization and long-term maintenance of potentiation, resulting in less reliable late-phase LTP.

**Figure 2.**
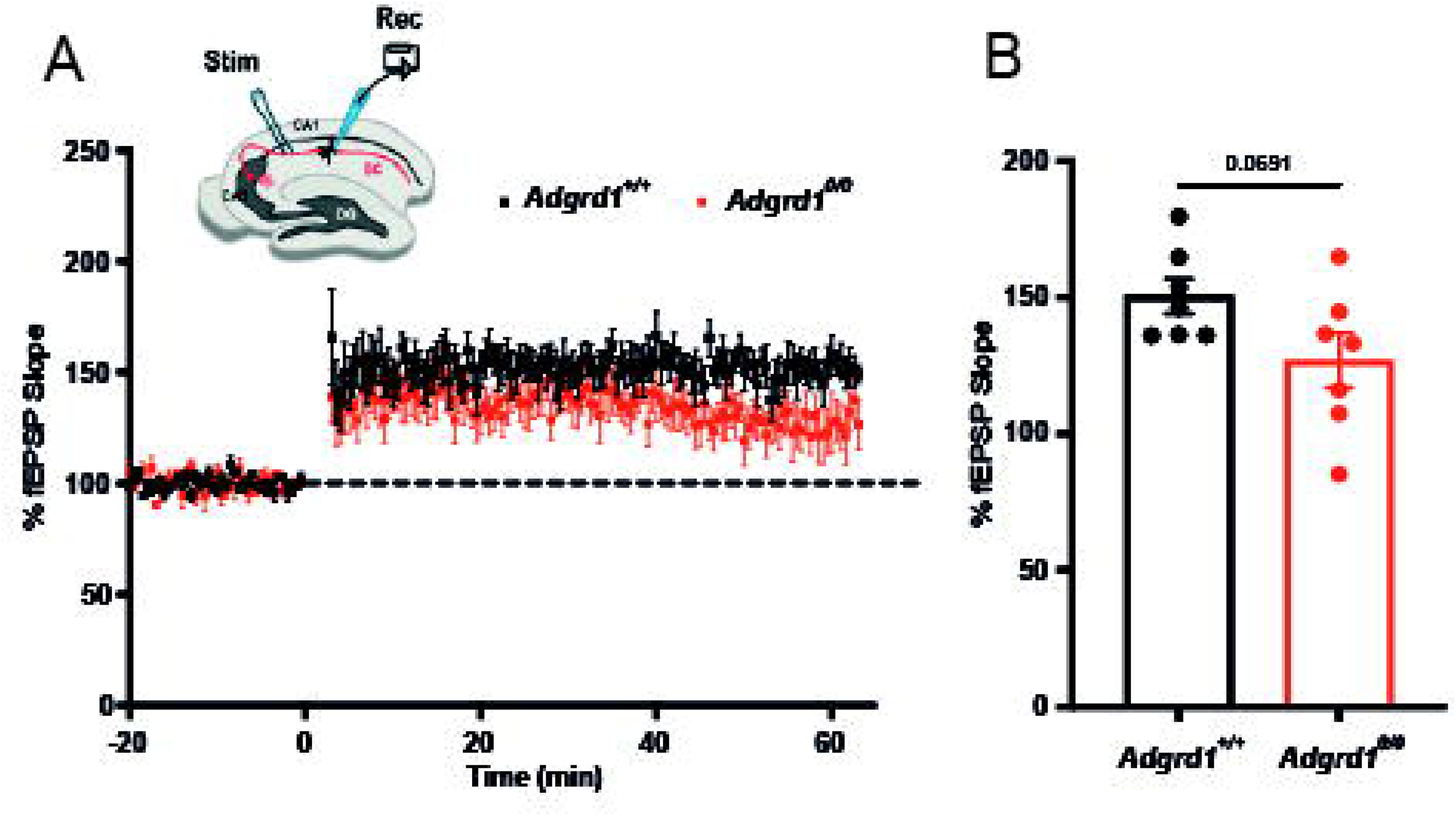
Electrophysiological assessment of synaptic plasticity in the hippocampal CA3-CA1 pathway. **A)** Time course of field excitatory postsynaptic potential (fEPSP) slope recorded in acute hippocampal slices following high-frequency stimulation. The schematic illustrates stimulation of Schaffer collateral (SC) fibers in CA3 and recording in the CA1 region. Baseline responses were comparable between genotypes; however, *Adgrd1^0/0^*slices exhibited a reduced and less stable potentiation compared to *Adgrd1^+/+^*controls during the maintenance phase of LTP. The dashed line indicates baseline levels (100%). **B)** Quantification of fEPSP slope during the late phase of LTP (indicated time window), showing a trend toward reduced potentiation in *Adgrd1^0/0^* mice relative to wild-type. Each point in B represents an individual slice. Statistical comparison was performed using an unpaired Student’s t-test.

### Enhanced susceptibility to KA in *Adgrd1* knockout mice

*Adgrd1* knockout and wild-type animals (C57BL/6J strain) were treated with KA administered intraperitoneally (Figure 3). After the last injection, the animals were recorded and monitored by two independent researchers for 4 hours (see Supplementary video 1 as example). Seizure severity was scored on a scale from I to VI. Results indicated that *Adgrd1^0/0^*mice exhibited a higher number of seizures and spent more time in advanced stages (grade VI) compared with wild-type animals. Specifically, the distribution of severity scores showed that mutant mice reached the following stages (Stage I-II: 56.25 %; Stage III-IV: 18.75 %; Stage V: 56.25 %; Stage VI: 81.25%; and Death: 25 %) and wild-type mice (Stage I-II: 54.92 %; Stage III-IV: 29.41 %; Stage V: 52.94 %; Stage VI: 23.53%; and Death: 5.88 %) (Figure 3A). Relevantly, an 81% and 25% of *Adgrd1^0/0^* reached Stage VI or died following the last KA treatment in contrast to wild-type mice. In addition, *Adgrd1^0/0^* mice showed increased seizure time (*Adgrd1^0/0^*: 69.30 ± 18.43 *vs.* wild-type: 2.79 ± 1.22; *p* = 0.0029, Mann-Whitney *U* test) (Figure 3B) with large number of seizures during the analysis (*Adgrd1^0^*^/*0*^: 12.88 ± 2.40 *vs.* wild-type: 5.70 ± 1.66; *p* = 0.030, Mann-Whitney *U* test) (Figure 3C).

**Figure 3.**
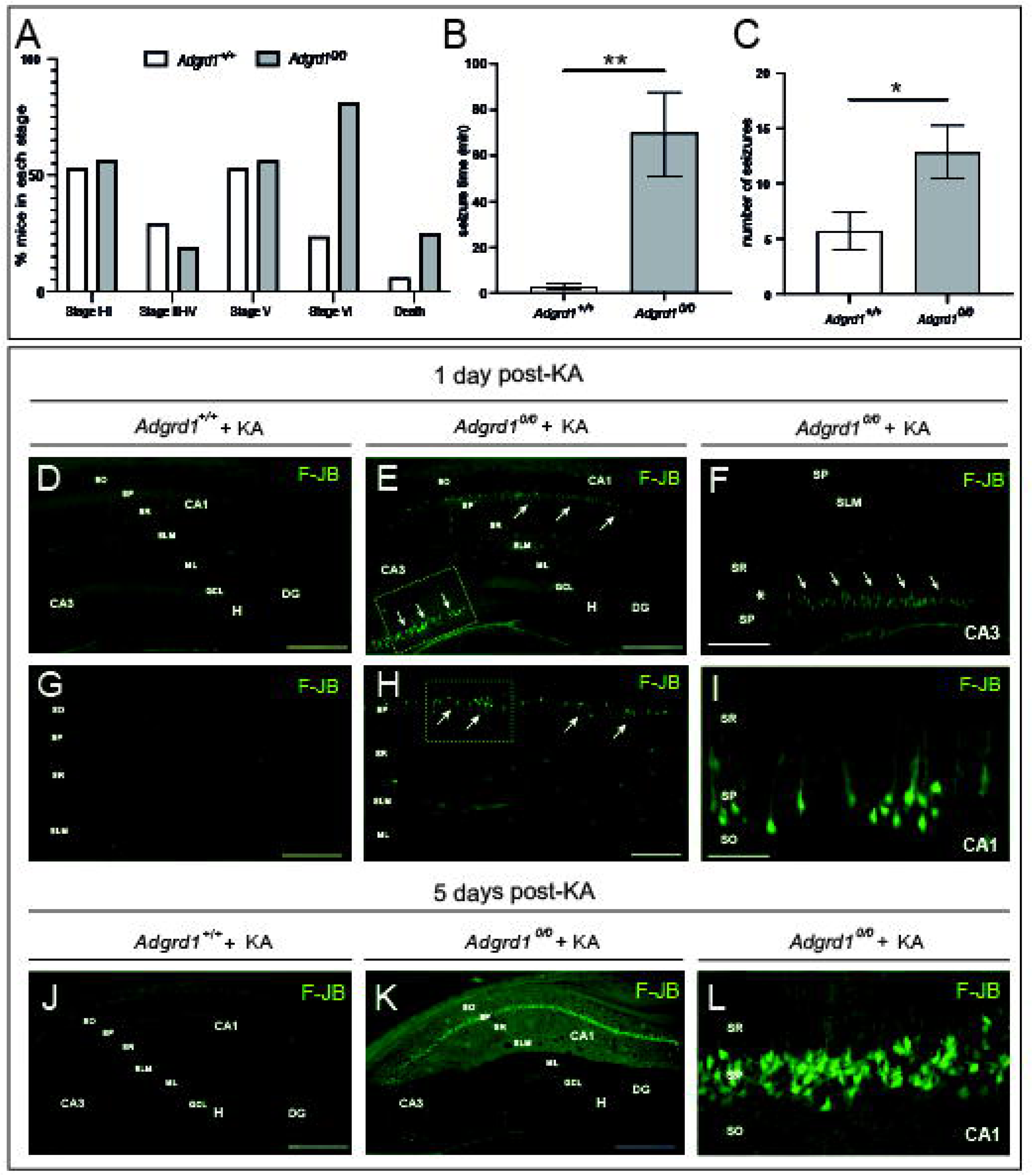
Enhanced KA susceptibility in Adrgd1 deficient mice. **A)** Percentage of mice reaching seizure stages I to VI and KA-induced mortality. Adult animals were subjected to a multiple KA-injection protocol (8 mg/kg b.w.) and epileptic responses were analyzed during 4 hours after the first KA injection **B, C)** Graphs illustrating the mean seizure time (B) and the total number of seizures (C) observed in the experiments between *Adgrd1^0/0^* and wild-type mice. Data is represented and mean ± SEM; *p* = 0.0029 and *p* = 0.030 Mann-Whitney *U* test in B and C respectively. **D-I)** Photomicrographs showing the pattern of neurodegeneration (Fluoro-Jade B staining) of hippocampus in *Adgrd1^+/+^*and *Adgrd1^0/0^* mice 1 day after KA treatment. Dying cells are mainly located in the pyramidal cell layer of CA1 and CA3 regions (arrows in E, F and H, I). F and I are high magnifications of the CA3 and CA1 regions showed in E and H respectively. **J-L)** Photomicrographs showing the pattern of Fluoro-Jade B staining of hippocampus in *Adgrd1^+/+^* and *Adgrd1^0/0^* mice 5 days after KA treatment. Increasing number of fluorescence cells can be observed in the CA1 of mutant mice (K, L) in contrast to wild-type (J). Abbreviations: CA1-3, *cornus ammonis 1- 3*; dg, *dentate gyrus*; gcl, granule cell layer; h, hilus; ml, molecular layer; sr, *stratum radiatum*; sl, *stratum lucidum*; slm, *stratum lacunosum-moleculare*; sp, *stratum pyramidale*; so, *stratum oriens*. Scale bars in D, E, G, H, J and K represents 200 µm. Scale bars in F, I and L represents 100 µm.

Once behavioral assessment following KA administration was completed, animals were processed for histological analysis either 1- or 5-days after the last KA injection (Figure 3D-L). Fluoro-Jade B staining revealed that, 1-day post-treatment, KA-treated *Adgrd1^0/0^* mice exhibited a high density of fluorescent cells in the pyramidal layer of the proximal CA3 region adjacent to the dentate gyrus, compared to wild-type animals (Figure 3D-F). Fewer labeled cells were observed in CA1, where fluorescent pyramidal neurons displayed a more dispersed distribution pattern (Figure 3G-I). Additionally, a small number of labeled cells was detected in the plexiform layers (Figure 3E, H). This spatial distribution closely resembled the pattern of Adgrd1 expression previously described by immunohistochemistry (Supplementary Figure 1).

At 5-days post-treatment, the number of labeled neurons in CA3 was markedly reduced, whereas the number of stained cells in the pyramidal layer of the CA1 was substantially increased (Figure 3K, L). This staining pattern was not observed in wild-type animals treated in parallel with KA (Figure 3J). Overall, these findings are consistent with the behavioral data, indicating that the pattern of neuronal degeneration closely parallels the severity and progression of seizures induced by KA.

### Enhanced astroglial and microglial proliferation and ERK1/2 kinase activity in Adgrd1 mice treated with KA

Intraperitoneal KA administration is well known to trigger a strong neuroinflammatory response in the hippocampus, characterized by pronounced astroglial and microglial activation and proliferation ^50,51^. After immunohistochemistry, we observed an increase in ERK1/2 levels in the mossy fibers of the hippocampus one day after KA administration compared with WT animals (Figure 4A, B). In addition, double immunostaining confirmed that the ERK1/2-positive cells observed only in the *Adgrd1^0/0^* hippocampus (Figure 4C) corresponded to astrocytes located in the CA1 and CA3 regions (Figure 4D-G). Similarly, after five days of treatment, *Adgrd1*-deficient animals showed a marked increase in activated microglia, identified by Iba1 immunostaining, in the hippocampus of KA-treated *Adgrd1^0/0^*mice compared with controls (Figure 4H- K).

**Figure 4.**
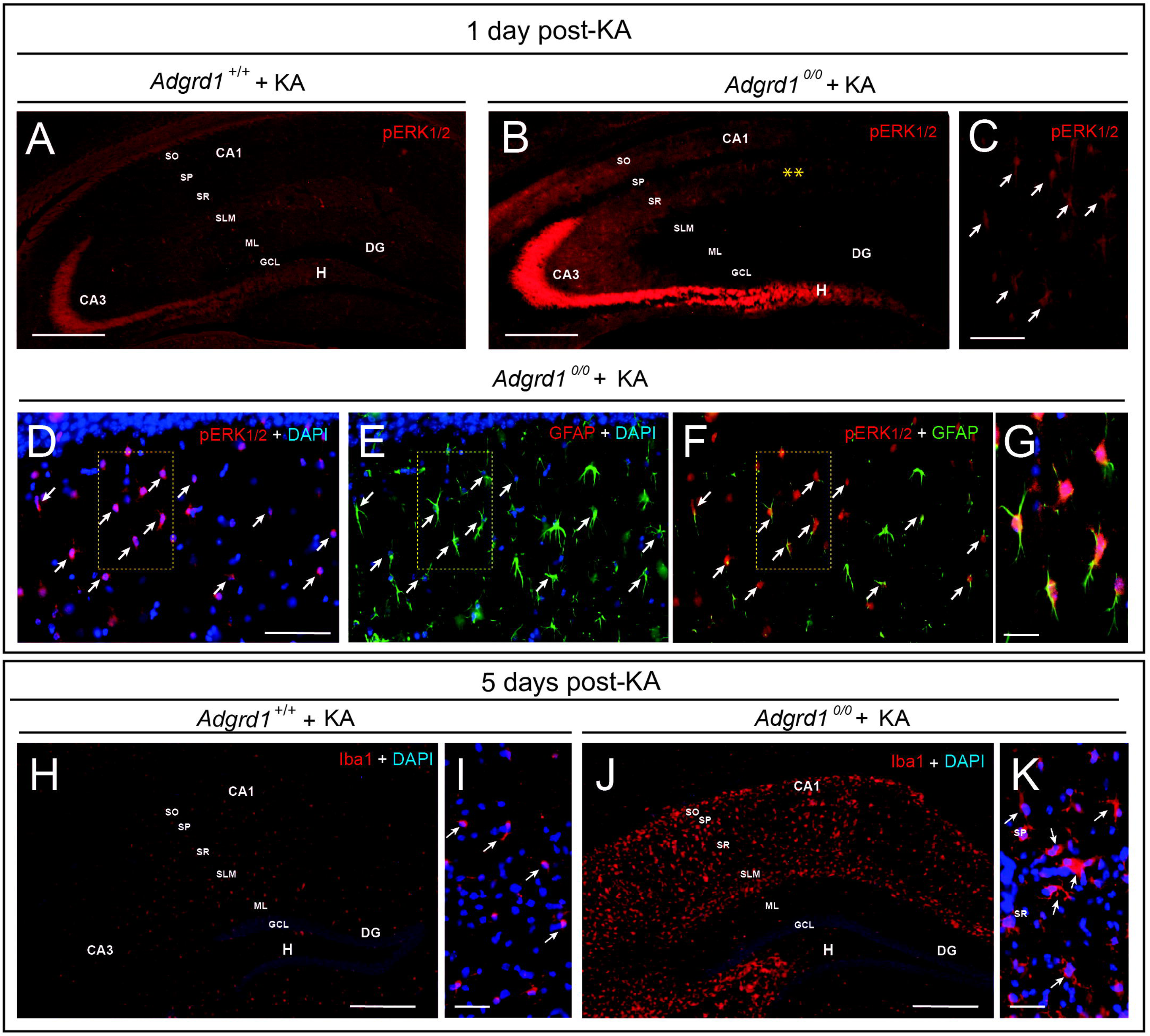
Increase gliosis and ERK1/2 activation in Adgrd1 deficient mice after KA treatment. **A-G)** Low (A, B) and medium (C-G) size fluorescence photomicrographs illustrating the increase of pERK1/2 one day after KA treatment in mutant mice. The mossy fibers and double labeled reactive astroglial cells (GFAP+ERK1/2) can be easily observed in CA1 and CA3 of treated *Adgrd1^0/0^* mice. **H-K)** Photomicrographs showing the increase in microglial cells 5-days after KA treatment in mutant compared to wild-type mice. Ameboid morphologies can be observed after Iba1 staining at higher magnification in treated *Adgrd1^0/0^* hippocampi. Abbreviations as in Figure 3. Scale bars in A, B, H and J represents 200 µm. Scale bars in C, D, G, I and K represents 100 µm.

### Transcriptomic alterations in the Adgrd-deficient hippocampus

To place the electrophysiological findings in a broader molecular context, we conducted a bulk RNA-seq analysis aimed at identifying transcriptional changes that could account for the alterations observed in hippocampal synaptic activity (Figure 5). Bulk RNA-seq analysis comparing adult hippocampal tissue from *Adgrd1^+/+^* and *Adgrd1^0/0^*mice identified a discrete set of differentially expressed genes (log2FC ≥ 1, p ≤ 0.05), indicating a targeted transcriptional response rather than widespread transcriptomic disruption (Figure 5). Consistent with this, principal component analysis revealed limited global separation between genotypes, suggesting that the transcriptional changes are driven by specific gene subsets rather than a dominant global shift (Figure 5A). Gene ontology enrichment analysis restricted to central nervous system (CNS)-related processes revealed a coordinated functional shift in *Adgrd1^0/0^* samples (Figure 5). Upregulated genes (n = 57) were primarily associated with neuronal signaling pathways, including G protein-coupled receptor signaling, cAMP-dependent pathways, calcium-mediated signaling, and positive regulation of neuron projection development. These functional categories are consistent with enhanced neuronal activity and synaptic signaling. At the gene level, this enrichment was driven by the upregulation of key neuronal regulators such as *Snca*, *Cox8b*, and *Dynlt1b*, which are associated with synaptic vesicle dynamics, mitochondrial function, and intracellular transport, respectively. In contrast, downregulated genes (n = 43) were enriched in processes related to ion transport, neurotransmitter transport, and inflammatory signaling pathways, including NF-κB and Toll-like receptor signaling. Notably, several genes involved in neuroinflammation and inhibitory neurotransmission, such as *Il1b*, *Gabra2*, and *Ltf*, were significantly reduced in *Adgrd1^0/0^* samples (Figure 5). Together, these results indicate that the absence of Adgrd1 induces a functional shift toward enhanced neuronal activity accompanied by a reduction in neuroinflammatory and inhibitory signaling, suggesting a potential imbalance in excitatory/inhibitory network regulation. Changes in selected genes were also measured and corroborated by RT-qPCR (Supplementary Figure 2).

**Figure 5.**
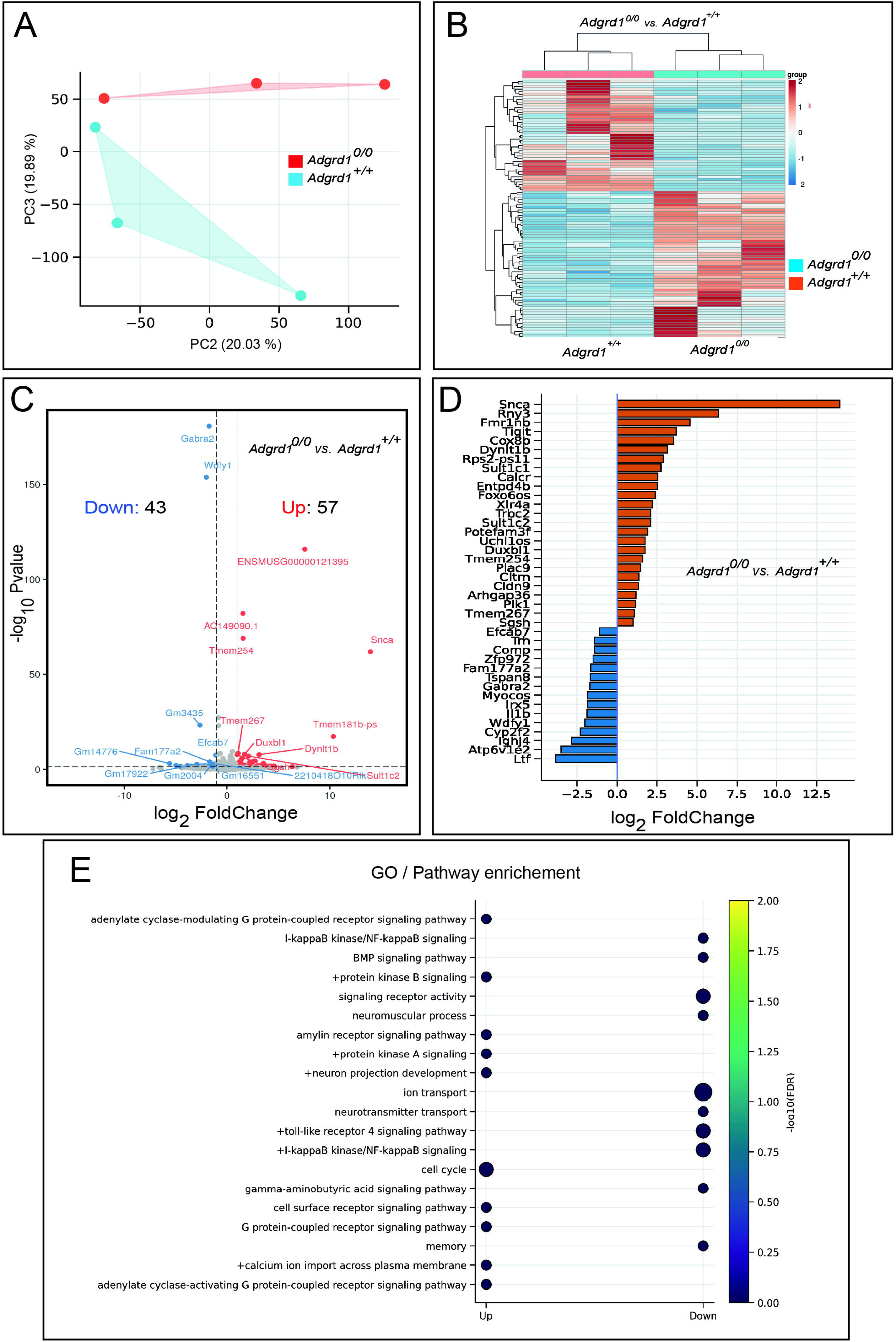
Transcriptomic alterations in the hippocampus of *Adgrd1^0/0^* mice. **A)** Principal component analysis (PCA) of RNA-seq samples shows only partial separation between *Adgrd1^0/0^* and *Adgrd1^+/+^*hippocampi, with variability largely captured by PC2 and PC3. This suggest that global transcriptomic differences between genotypes are modest and not driven by a dominant shift across the entire transcriptome. **B)** Unsupervised hierarchical clustering of differentially expressed genes reveals a clear segregation of samples according to genotype, indicating the presence of a consistent transcriptional signature distinguishing *Adgrd1^0/0^* from *Adgrd1^+/+^* hippocampi. **C)** Volcano plot summarizing differential gene expression (log₂FC ≥ 1, p ≤ 0.05) identifies 57 upregulated and 43 downregulated genes in *Adgrd1^0/0^* samples *vs. Adgrd1^+/+^*. Notable downregulated genes include *Gabra2* and *Wdfy1*, whereas *Snca*, *Dynlt1b*, and *Duxbl1* are among the most prominently upregulated transcripts. **D)** Barplot of selected annotated genes highlights the magnitude and direction of expression changes, emphasizing strong upregulation of genes such as *Snca* and more moderate but consistent downregulation of genes including *Gabra2*, *Il1b*, and *Wdfy1***. E)** Gene ontology (GO) enrichment analysis focused on CNS-related biological processes is represented as a dot heatmap, where point size corresponds to the number of genes associated with each GO term and color indicates statistical significance (−log10 FDR). Enrichment analysis was performed using a hypergeometric test with multiple testing correction. Notice the down regulation in GABA-mediated signaling and memory pathways, as well as other pathways (e.g., ion transport, signaling receptor activity, etc).

## Discussion

### The absence of Adgrd1 disrupts hippocampal resilience and circuit stability

Our findings identify *Adgrd1* as a regulator of hippocampal robustness, influencing both neuronal survival and circuit-level function. *Adgrd1^0/0^*mice exhibited heightened susceptibility to KA-induced excitotoxicity, with pronounced neuronal degeneration in CA3 and CA1 accompanied by astroglia and microglia activation as well established ^52^. This phenotype suggests that Adrgrd1 contributes to mechanisms that buffer hippocampal networks against hyperexcitability and metabolic stress, consistent with the emerging role of adhesion GPCRs in maintaining neuronal integrity, mechanotransduction, and extracellular communication (see Introduction for references). Although this vulnerability parallels that observed in *Prnp^ZH3/ZH3^*mice ^26^, the behavior and underlying molecular signatures diverge substantially (see below), indicating distinct resilience pathways.

### Behavioral alterations suggest impaired exploratory drive and disrupted hippocampal-prefrontal interactions

The hippocampal-prefrontal circuit supports the integration of spatial, mnemonic, and executive processes, enabling flexible, goal-directed behavior based on prior experience and contextual information ^53–55^. Behaviorally, *Adgrd1^0/0^*mice exhibited impaired nest building, reduced object exploration, and increased immobility in the center of the open-field arena. This behavioral profile does not align with classical anxiolytic or anxiogenic phenotypes ^56,57^, nor does it indicate overt motor deficits ^58^, but rather suggests reduced exploratory engagement and impaired initiation of goal-directed actions. Consistently, while recognition memory appeared preserved, as indicated by an intact discrimination index in the NOR task, decreased object interaction further supports a reduction in motivational drive. Increased immobility in the center zone, an area typically avoided, also points to altered spatial evaluation and action initiation ^59–61^, a behavioral pattern associated with imbalances in hippocampal excitation/inhibition dynamics ^62–64^, in line with the transcriptional and electrophysiological alterations observed in this study.

### Electrophysiological evidence reveals intact basal transmission but impaired LTP maintenance

Electrophysiological recordings demonstrated that loss of Adrgd1 function does not affect basal synaptic transmission: input-output curves and paired-pulse facilitation were indistinguishable between genotypes, indicating preserved presynaptic release probability ^65^. LTP induction was also normal, but late-phase maintenance was altered, with *Adgrd1^0/0^* slices showing reduced potentiation amplitude and increased variability. Given that Adgrd1 couples predominantly to G_s_ and modulate intracellular cAMP, its absence may weaken signaling pathways that rely on cAMP-dependent activation, specially CREB. This pathway is a well-established molecular axis required for the stabilization of late-phase LTP _66,67_. Importantly, impairments in the cAMP-CREB cascade selectively disrupt LTP maintenance while sparing its induction ^66,68^, a pattern that closely resembles the electrophysiological profile observed in *Adgrd1^0/0^* slices.

### Transcriptomic alterations provide a mechanistic substrate for impaired LTP stabilization

RNA-seq analysis revealed a selective transcriptional signature in *Adgrd1^0/0^* hippocampi. Downregulated genes included *Gabra2, Wdfy1, Irx5, Comp, Efcab7, Il1b,* and *Trh*, implicating deficits in inhibitory signaling, extracellular matrix organization, calcium-dependent processes, and neuroimmune modulation; pathways essential for synaptic stabilization and the maintenance of excitation/inhibition balance ^62^. Conversely, upregulated genes such as *Snca, Arhgap36, Dynlt1b, Calcr, Sgsh, Entpd4b, Uchl1os,* and immune-related transcripts (*Tigit, Trac* and *Trbc2*) suggest enhanced presynaptic remodeling, altered GPCR signaling, cytoskeletal reorganization, and a more reactive cellular environment (e.g., ^69–73^). Together, this shift, characterized by the loss of synaptic stabilizing components alongside the emergence of remodeling and inflammatory signatures, provides a coherent molecular framework for the observed electrophysiological phenotype. In particular, reduced *Gabra2* expression may weaken inhibitory tone (e.g., ^74^), while alterations in extracellular matrix and calcium-associated pathways may compromise structural consolidation (e.g., ^75^). In parallel, the upregulation of *Snca* and cytoskeletal regulators is consistent with compensatory presynaptic adaptations that may, however, destabilize long-term plasticity ^76–78^. Although some of the affected genes are linked to activity-dependent transcriptional programs, our data do not support a direct or selective impairment of a single upstream transcriptional regulator but rather point to a broader disruption of coordinated gene networks underlying hippocampal circuit function.

### Divergence from *Prnp* deletion models highlights a distinct functional role **for *Adgrd1***

Although both *Adgrd1^0/0^* and *Prnp^ZH3/ZH3^* mice exhibit increased susceptibility to KA-induced seizures, the molecular architecture underlying this vulnerability differs substantially between the two models. In *Prnp*-deficient mice, reduced expression of the GABA_A_ receptor α2 subunit constitutes a primary mechanism driving hippocampal hyperexcitability, consistent with previous work demonstrating impaired inhibitory homeostasis and heightened sensitivity to strong stimulation ^26^. In contrast, while *Adgrd1^0/0^* mice also show downregulation of *Gabra2*, this occurs within a broader transcriptional landscape involving cytoskeletal regulators, extracellular matrix components, immune-related genes, and presynaptic stress markers such as *Snca*. This pattern suggests that the reduction GABA_A_R α2 in *Adgrd1^0/0^* mice is unlikely to reflect a direct mechanistic parallel to PrP^C^ loss, but rather a downstream consequence of altered cAMP-dependent signaling, impaired synaptic stabilization, or increased hippocampal stress sensitivity. Consistent with this interpretation, hippocampal *Prnp* expression shows a modest, non-significant increase in *Adgrd1^0/0^* mice, that might reflect a stress-responsive adaptation rather than a primary molecular driver (e.g., ^79^). Together, these observations indicate that the two systems may converge on a similar physiological phenotype through distinct upstream mechanisms.

Taken together, the behavioral, electrophysiological and transcriptomic data support a model in which Adgrd1 contributes to sustaining the molecular framework required for long-term synaptic stability and coherent exploratory behavior. Although basal synaptic transmission appears largely unaffected by its deletion, the absence of Adgrd1 alters gene expression programs linked to the consolidation phase of synaptic plasticity. This is in line with the observed deficits in LTP maintenance and the reduction in exploratory activity. In addition, these changes may render the system more susceptible to excitotoxic conditions. Further studies will be necessary to clarify the underlying cellular and molecular mechanisms, but the present findings point to Adgrd1 as a relevant component in maintaining hippocampal circuit stability and adaptive behavioral responses in young-adult mice.

## Data Availability Statement

The raw data supporting the conclusions of this article will be made available by the authors upon request, without undue reservation.

## Author Contributions

IMS, PPP, APG, NM, KZ performed experiments PPP, EDC, AA, JD, XG, RG and JADR designed experiments, JADR, IMS and PPP edited the manuscript. JADR provided experimental funds. All authors contributed to the article and approved the submitted version.

## Acknowledgements.

The authors thank Miriam Segura-Feliu and Margarita Carmona for their technical help. We also thank the microFab nanotechnology platform of IBEC for their technical help and the Genomic Unit from Scientific and Technological Centers (CCiTUB), Universitat de Barcelona. The authors also thank all the members of the J.A. del Río, A. Aguzzís laboratories for their helpful comments and support.

---

**Supplementary Figure 1. Genotyping of *Adgrd1* mice and histological characterization of Adgrd1 expression in wild-type hippocampus.**

**A)** Representative PCR genotyping showing the identification of *Adgrd1^+/+^*, *Adgrd1^+/−^*, and *Adgrd1^0/0^* alleles, with bands at approximately 295 bp and 170 bp as indicated. Molecular weight markers are shown on both sides.

**B)** Immunohistochemical staining in kidney sections showing sparse positive interstitial cells (arrows) immunopositive for Adgrd1. **C)** Low-magnification photomicrograph illustrating immunolabeling with the α-Adgrd1 antibody in the adult hippocampus after nickel-enhanced DAB development, with immunopositive cells observed in the *stratum pyramidale* of the CA3 region. **D, E)** Representative high-magnification images of hippocampal regions (CA3 and associated layers) showing labeled cells (arrows) in wild-type mice; note that not all pyramidal neurons in CA3 are labeled (arrowheads in D). After nickel enhancement, immunopositive cells located in plexiform layers become more visible, although accompanied by increased background staining. **F, G)** Low- magnification hippocampal sections illustrating background staining controls, including (F) *Adgrd1^0/0^*sections incubated with the α-Adgrd1 antibody and developed with prolonged peroxidase reaction, and (G) wild-type sections processed with omission of the primary antibody. Abbreviations in C-G as in Figure 3. UC, ureteric collecting duct; KG, kidney glomerulus. Scale bars: B, C, F and G = 200 µm; D, E = 100 µm.

**Supplementary Figure 2. RT-qPCR analysis of gene expression in Adgrd1 mutant mice.**

**A)** Relative mRNA expression levels (2^−ΔCt^) of the indicated genes in *Adgrd1^+/+^*(white bars) and *Adgrd1^0/0^* (grey bars) samples. Data are presented as mean ± SEM, with individual data points overlaid. Dashed lines indicate reference expression thresholds. Statistically significant differences between genotypes are indicated above the corresponding comparisons (*p* values shown).

## References

1 Vizurraga, A., Adhikari, R., Yeung, J., Yu, M. & Tall, G. G. Mechanisms of adhesion G protein-coupled receptor activation. J Biol Chem 295, 14065– 14083, doi:10.1074/jbc.REV120.007423 (2020).

2 Bohnekamp, J. & Schöneberg, T. Cell adhesion receptor GPR133 couples to Gs protein. J Biol Chem 286, 41912–41916, doi:10.1074/jbc.C111.265934 (2011).

3 Lala, T. & Hall, R. A. Adhesion G protein-coupled receptors: structure, signaling, physiology, and pathophysiology. Physiological Reviews 102, 1587–1624, doi:10.1152/physrev.00027.2021 (2022).

4 Hamann, J. et al. International Union of Basic and Clinical Pharmacology. XCIV. Adhesion G Protein-Coupled Receptors. Pharmacological Reviews 67, 338–367, doi:10.1124/pr.114.009647 (2015).

5 Fischer, L., Wilde, C., Schöneberg, T. & Liebscher, I. Functional relevance of naturally occurring mutations in adhesion G protein-coupled receptor ADGRD1 (GPR133). BMC Genomics 17, 609, doi:10.1186/s12864-016-2937-2 (2016).

6 Bjarnadóttir, T. K. et al. The human and mouse repertoire of the adhesion family of G-protein-coupled receptors. Genomics 84, 23–33, 10.1016/j.ygeno.2003.12.004 (2004).

7 Bjarnadóttir, T. K. et al. Identification of novel splice variants of Adhesion G protein-coupled receptors. Gene 387, 38–48, doi:10.1016/j.gene.2006.07.039 (2007).

8 Araç, D. et al. A novel evolutionarily conserved domain of cell-adhesion GPCRs mediates autoproteolysis. Embo j 31, 1364–1378, doi:10.1038/emboj.2012.26 (2012).

9 Strokes, N. & Piao, X. in Adhesion-GPCRs: Structure to Function (eds Simon Yona & Martin Stacey) 87–97 (Springer US, 2010).

10 Bayin, N. S. et al. GPR133 (ADGRD1), an adhesion G-protein-coupled receptor, is necessary for glioblastoma growth. Oncogenesis 5, e263, doi:10.1038/oncsis.2016.63 (2016).

11 Frenster, J. D. et al. Expression profiling of the adhesion G protein-coupled receptor GPR133 (ADGRD1) in glioma subtypes. Neurooncol Adv 2, vdaa053, doi:10.1093/noajnl/vdaa053 (2020).

12 Yang, Z. et al. Identification, structure, and agonist design of an androgen membrane receptor. Cell 188, 1589–1604.e1524, 10.1016/j.cell.2025.01.006 (2025).

13 Lehmann, J. et al. The mechanosensitive adhesion G protein-coupled receptor 133 (GPR133/ADGRD1) enhances bone formation. Signal Transduction and Targeted Therapy 10, 199, doi:10.1038/s41392-025-02291-y (2025).

14 Stephan, G. et al. Modulation of GPR133 (ADGRD1) signaling by its intracellular interaction partner extended synaptotagmin 1. Cell Rep 43, 114229, doi:10.1016/j.celrep.2024.114229 (2024).

15. He, L., et al. Exogenous activation of the adhesion GPCR ADGRD1/GPR133 protects against bone loss by negatively regulating osteoclastogenesis. (2025).

16 Küffer, A. et al. The prion protein is an agonistic ligand of the G protein-coupled receptor Adgrg6. Nature 536, 464–468, doi:10.1038/nature19312 (2016).

17 Kovač, V. & Čurin Šerbec, V. Prion Protein: The Molecule of Many Forms and Faces. Int J Mol Sci 23, 1232 (2022).

18 Cha, S. & Kim, M. Y. The role of cellular prion protein in immune system. BMB Rep 56, 645–650, doi:10.5483/BMBRep.2023-0151 (2023).

19 Scialo, C. & Legname, G. The role of the cellular prion protein in the uptake and toxic signaling of pathological neurodegenerative aggregates. Prog Mol Biol Transl Sci 175, 297–323, doi:10.1016/bs.pmbts.2020.08.008 (2020).

20 Watts, J. C., Bourkas, M. E. C. & Arshad, H. The function of the cellular prion protein in health and disease. Acta Neuropathol 135, 159–178, doi:10.1007/s00401-017-1790-y (2018).

21 Del Rio, J. A., Ferrer, I. & Gavin, R. Role of cellular prion protein in interneuronal amyloid transmission. Prog Neurobiol 165-167, 87–102, doi:10.1016/j.pneurobio.2018.03.001 (2018).

22 Walz, R. et al. Cellular prion protein: implications in seizures and epilepsy. Cell Mol Neurobiol 22, 249–257, doi:10.1023/a:1020711700048 (2002).

23 Linden, R. The Biological Function of the Prion Protein: A Cell Surface Scaffold of Signaling Modules. Frontiers in Molecular Neuroscience Volume 10, doi:10.3389/fnmol.2017.00077 (2017).

24 Westergard, L., Christensen, H. M. & Harris, D. A. The cellular prion protein (PrP(C)): its physiological function and role in disease. Biochim Biophys Acta 1772, 629–644, doi:10.1016/j.bbadis.2007.02.011 (2007).

25 Chrobak, A. A. et al. New Light on Prions: Putative Role of PrPc in Pathophysiology of Mood Disorders. International Journal of Molecular Sciences 25, 2967 (2024).

26 Matamoros-Angles, A. et al. Analysis of co-isogenic prion protein deficient mice reveals behavioral deficits, learning impairment, and enhanced hippocampal excitability. BMC Biol 20, 17, doi:10.1186/s12915-021-01203-0 (2022).

27 Gavin, R. & Del Rio, J. A. Exploring the Biological Connection Between Tau and PrP(C) in Neuronal Cells: GSK3beta as a Possible Key Player. Mol Neurobiol 62, 15284–15294, doi:10.1007/s12035-025-05163-2 (2025).

28 Coitinho, A. S. et al. The interaction between prion protein and laminin modulates memory consolidation. Eur J Neurosci 24, 3255–3264, doi:10.1111/j.1460-9568.2006.05156.x (2006).

29 Chrobak, A. A. et al. New Light on Prions: Putative Role of PrP(c) in Pathophysiology of Mood Disorders. Int J Mol Sci 25, doi:10.3390/ijms25052967 (2024).

30 Gadotti, V. M., Bonfield, S. P. & Zamponi, G. W. Depressive-like behaviour of mice lacking cellular prion protein. Behav Brain Res 227, 319–323, doi:10.1016/j.bbr.2011.03.012 (2012).

31 Carulla, P. et al. Involvement of PrP(C) in kainate-induced excitotoxicity in several mouse strains. Sci Rep 5, 11971, doi:10.1038/srep11971 (2015).

32 Llorens, F. & Del Rio, J. A. Unraveling the neuroprotective mechanisms of PrP (C) in excitotoxicity. Prion 6, 245–251, doi:10.4161/pri.19639 (2012).

33 Carulla, P. et al. Neuroprotective role of PrPC against kainate-induced epileptic seizures and cell death depends on the modulation of JNK3 activation by GluR6/7-PSD-95 binding. Mol Biol Cell 22, 3041–3054, doi:10.1091/mbc.E11-04-0321 (2011).

34 Rangel, A. et al. Regulation of GABA(A) and glutamate receptor expression, synaptic facilitation and long-term potentiation in the hippocampus of prion mutant mice. PLoS One 4, e7592, doi:10.1371/journal.pone.0007592 (2009).

35 Rangel, A. et al. Enhanced susceptibility of Prnp-deficient mice to kainate-induced seizures, neuronal apoptosis, and death: Role of AMPA/kainate receptors. J Neurosci Res 85, 2741–2755, doi:10.1002/jnr.21215 (2007).

36 Striebel, J. F., Race, B. & Chesebro, B. Prion protein and susceptibility to kainate-induced seizures: genetic pitfalls in the use of PrP knockout mice. Prion 7, 280–285, doi:10.4161/pri.25738 (2013).

37 Walz, R. et al. Increased sensitivity to seizures in mice lacking cellular prion protein. Epilepsia 40, 1679–1682, doi:10.1111/j.1528-1157.1999.tb01583.x (1999).

38 Striebel, J. F., Race, B., Pathmajeyan, M., Rangel, A. & Chesebro, B. Lack of influence of prion protein gene expression on kainate-induced seizures in mice: studies using congenic, coisogenic and transgenic strains. Neuroscience 238, 11–18, doi:10.1016/j.neuroscience.2013.02.004 (2013).

39 Deacon, R. M. Assessing nest building in mice. Nat Protoc 1, 1117–1119, doi:10.1038/nprot.2006.170 (2006).

40 Miedel, C. J., Patton, J. M., Miedel, A. N., Miedel, E. S. & Levenson, J. M. Assessment of Spontaneous Alternation, Novel Object Recognition and Limb Clasping in Transgenic Mouse Models of Amyloid-β and Tau Neuropathology. J Vis Exp, doi:10.3791/55523 (2017).

41 Bello-Arroyo, E. et al. MouBeAT: A New and Open Toolbox for Guided Analysis of Behavioral Tests in Mice. Front Behav Neurosci 12, 201, doi:10.3389/fnbeh.2018.00201 (2018).

42 Seibenhener, M. L. & Wooten, M. C. Use of the Open Field Maze to measure locomotor and anxiety-like behavior in mice. J Vis Exp, e52434, doi:10.3791/52434 (2015).

43 Soriano, E. & Del Rio, J. A. Simultaneous immunocytochemical visualization of bromodeoxyuridine and neural tissue antigens. J Histochem Cytochem 39, 255–263, doi:10.1177/39.3.1671576 (1991).

44 Adams, J. C. Heavy metal intensification of DAB-based HRP reaction product. J Histochem Cytochem 29, 775, doi:10.1177/29.6.7252134 (1981).

45 Chen, S., Zhou, Y., Chen, Y. & Gu, J. fastp: an ultra-fast all-in-one FASTQ preprocessor. Bioinformatics 34, i884–i890, doi:10.1093/bioinformatics/bty560 (2018).

46 Kim, D., Langmead, B. & Salzberg, S. L. HISAT: a fast spliced aligner with low memory requirements. Nat Methods 12, 357–360, doi:10.1038/nmeth.3317 (2015).

47 Li, H. et al. The Sequence Alignment/Map format and SAMtools. Bioinformatics 25, 2078–2079, doi:10.1093/bioinformatics/btp352 (2009).

48 Anders, S., Pyl, P. T. & Huber, W. HTSeq--a Python framework to work with high-throughput sequencing data. Bioinformatics 31, 166–169, doi:10.1093/bioinformatics/btu638 (2015).

49 Love, M. I., Huber, W. & Anders, S. Moderated estimation of fold change and dispersion for RNA-seq data with DESeq2. Genome Biol 15, 550, doi:10.1186/s13059-014-0550-8 (2014).

50 Vezzani, A. & Granata, T. Brain inflammation in epilepsy: experimental and clinical evidence. Epilepsia 46, 1724–1743, doi:10.1111/j.1528-1167.2005.00298.x (2005).

51 Vezzani, A., French, J., Bartfai, T. & Baram, T. Z. The role of inflammation in epilepsy. Nat Rev Neurol 7, 31–40, doi:10.1038/nrneurol.2010.178 (2011).

52 Ben-Ari, Y. Limbic seizure and brain damage produced by kainic acid: Mechanisms and relevance to human temporal lobe epilepsy. Neuroscience 14, 375–403, doi:10.1016/0306-4522(85)90299-4 (1985).

53 Eichenbaum, H. Prefrontal–hippocampal interactions in episodic memory. Nature Reviews Neuroscience 18, 547–558, doi:10.1038/nrn.2017.74 (2017).

54 Sigurdsson, T. & Duvarci, S. Hippocampal-Prefrontal Interactions in Cognition, Behavior and Psychiatric Disease. Frontiers in Systems Neuroscience **Volume** 9 - 2015, doi:10.3389/fnsys.2015.00190 (2016).

55 Jin, J. & Maren, S. Prefrontal-Hippocampal Interactions in Memory and Emotion. Frontiers in Systems Neuroscience **Volume** 9 - 2015, doi:10.3389/fnsys.2015.00170 (2015).

56 Lezak, K. R., Missig, G. & Carlezon, W. A., Jr. Behavioral methods to study anxiety in rodents. Dialogues Clin Neurosci 19, 181–191, doi:10.31887/DCNS.2017.19.2/wcarlezon (2017).

57 Heinz, D. E. et al. Exploratory drive, fear, and anxiety are dissociable and independent components in foraging mice. Transl Psychiatry 11, 318 10.1038/s41398-021-01458-9 (2021).

58 Demchuk, A. M., Esteves, I. M. & McNaughton, B. L. Non-maternal nest building behaviours in mice predict bilateral dorsal hippocampal lesion extent. Behav Brain Res 480, 115366, doi:10.1016/j.bbr.2024.115366 (2025).

59 Cohen, S. J. & Stackman, R. W., Jr. Assessing rodent hippocampal involvement in the novel object recognition task. A review. Behav Brain Res 285, 105–117, doi:10.1016/j.bbr.2014.08.002 (2015).

60 Figueiredo Cerqueira, M. M., et al. Comparative analysis between Open Field and Elevated Plus Maze tests as a method for evaluating anxiety-like behavior in mice. Heliyon 9, e14522, doi:10.1016/j.heliyon.2023.e14522 (2023).

61 Kay, K. et al. A hippocampal network for spatial coding during immobility and sleep. Nature 531, 185–190, doi:10.1038/nature17144 (2016).

62 Bannerman, D. M. et al. Hippocampal synaptic plasticity, spatial memory and anxiety. Nat Rev Neurosci 15, 181–192, doi:10.1038/nrn3677 (2014).

63 Malagon-Vina, H., Ciocchi, S. & Klausberger, T. Firing patterns of ventral hippocampal neurons predict the exploration of anxiogenic locations. eLife 12, e83012, doi:10.7554/eLife.83012 (2023).

64 Bonansco, C. & Fuenzalida, M. Plasticity of Hippocampal Excitatory-Inhibitory Balance: Missing the Synaptic Control in the Epileptic Brain. Neural Plast 2016, 8607038, doi:10.1155/2016/8607038 (2016).

65 Jackman, S. L. & Regehr, W. G. The Mechanisms and Functions of Synaptic Facilitation. Neuron 94, 447–464, doi:10.1016/j.neuron.2017.02.047 (2017).

66 Abel, T. et al. Genetic demonstration of a role for PKA in the late phase of LTP and in hippocampus-based long-term memory. Cell 88, 615–626, doi:10.1016/s0092-8674(00)81904-2 (1997).

67 Barco, A., Alarcon, J. M. & Kandel, E. R. Expression of constitutively active CREB protein facilitates the late phase of long-term potentiation by enhancing synaptic capture. Cell 108, 689–703,10.1016/s0092-8674(02)00657-8 (2002).

68 Kida, S. A Functional Role for CREB as a Positive Regulator of Memory Formation and LTP. Exp Neurobiol 21, 136–140, doi:10.5607/en.2012.21.4.136 (2012).

69 Sharma, M. & Burré, J. α-Synuclein in synaptic function and dysfunction. Trends Neurosci 46, 153–166, doi:10.1016/j.tins.2022.11.007 (2023).

70 Burré, J. The Synaptic Function of α-Synuclein. J Parkinsons Dis 5, 699– 713, doi:10.3233/jpd-150642 (2015).

71 Ostrovskaya, A. et al. Expression and activity of the calcitonin receptor family in a sample of primary human high-grade gliomas. BMC Cancer 19, 157, doi:10.1186/s12885-019-5369-y (2019).

72 Hirokawa, N., Niwa, S. & Tanaka, Y. Molecular Motors in Neurons: Transport Mechanisms and Roles in Brain Function, Development, and Disease. Neuron 68, 610–638, doi:10.1016/j.neuron.2010.09.039 (2010).

73 Hashimoto-Tane, A. et al. Dynein-Driven Transport of T Cell Receptor Microclusters Regulates Immune Synapse Formation and T Cell Activation. Immunity 34, 919–931, doi:10.1016/j.immuni.2011.05.012 (2011).

74 Yu, W. et al. Gabra2 is a genetic modifier of Scn8a encephalopathy in the mouse*. Epilepsia 61, 2847–2856, 10.1111/epi.16741 (2020).

75 Yang, L., Wei, M., Xing, B. & Zhang, C. Extracellular matrix and synapse formation. Bioscience Reports 43, doi:10.1042/bsr20212411 (2023).

76 Jędrzejewska-Szmek, J. & Blackwell, K. T. From membrane receptors to protein synthesis and actin cytoskeleton: Mechanisms underlying long lasting forms of synaptic plasticity. Semin Cell Dev Biol 95, 120–129, doi:10.1016/j.semcdb.2019.01.006 (2019).

77 Ramalingam, N. et al. Dynamic physiological α-synuclein S129 phosphorylation is driven by neuronal activity. npj Parkinson’s Disease 9, 4, doi:10.1038/s41531-023-00444-w (2023).

78 Nemani, V. M. et al. Increased expression of alpha-synuclein reduces neurotransmitter release by inhibiting synaptic vesicle reclustering after endocytosis. Neuron 65, 66–79, doi:10.1016/j.neuron.2009.12.023 (2010).

79 Zeng, L., Zou, W. & Wang, G. Cellular prion protein (PrP(C)) and its role in stress responses. Int J Clin Exp Med 8, 8042–8050 (2015).

