## Supplementary figures and images for "Adgrd1 deficiency reveals increased hippocampal vulnerability and selective behavioral alterations in mice"

### Supp Figure 2

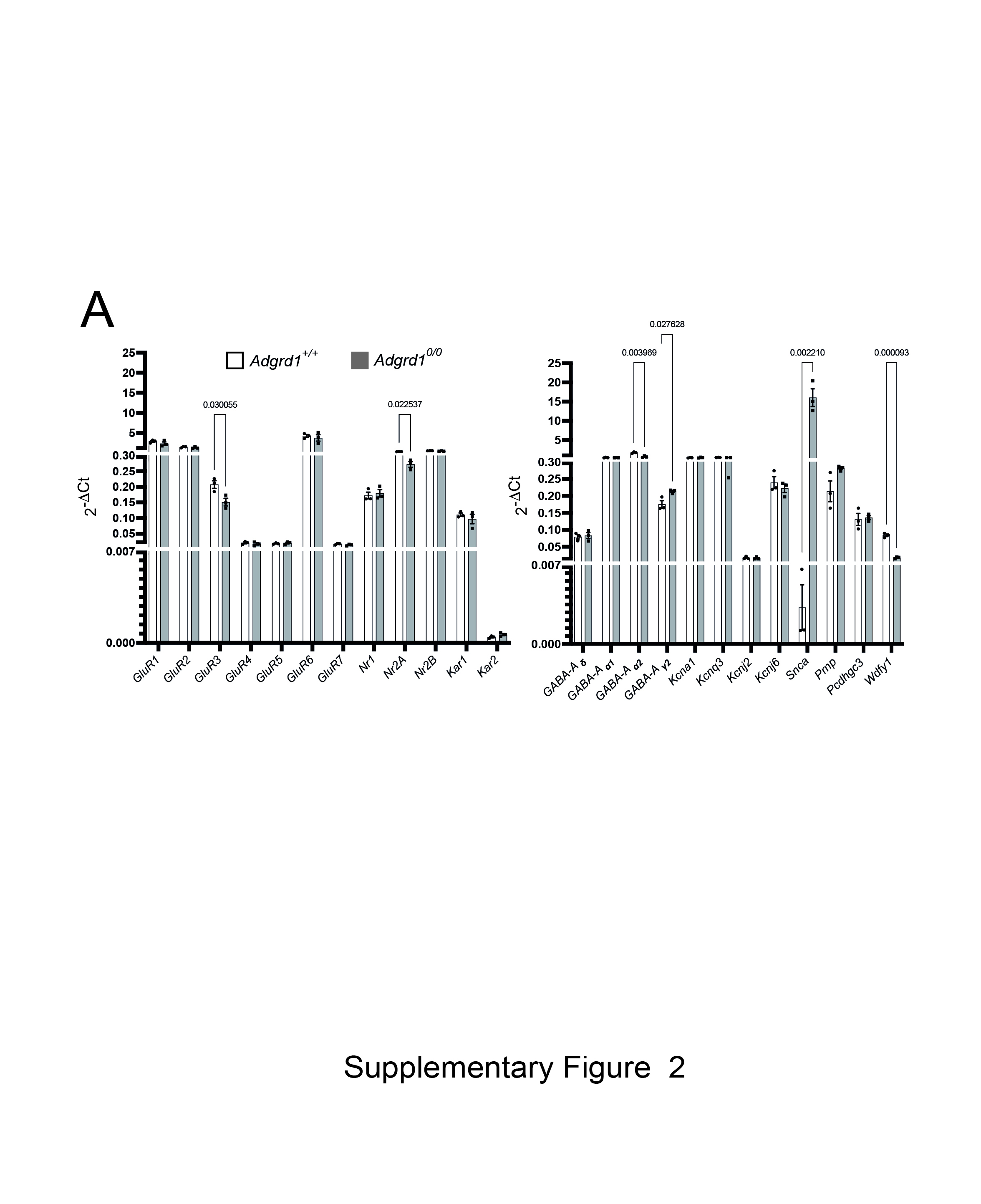

### Suppl Figure 1

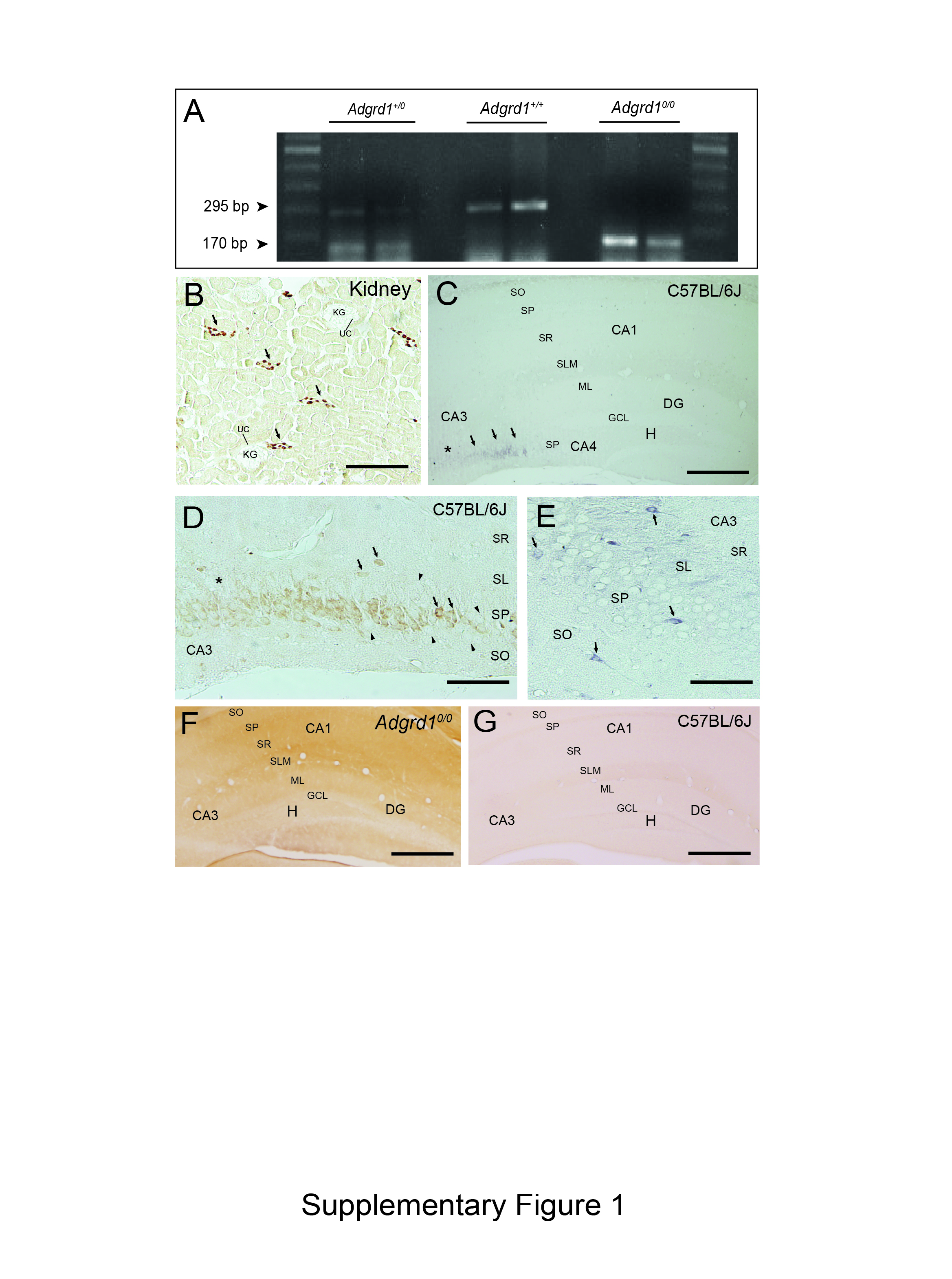
